# RNA strand invasion activity of the Polycomb complex PRC2

**DOI:** 10.1101/635722

**Authors:** Célia Alecki, Victoria Chiwara, Lionel A. Sanz, Daniel Grau, Osvaldo Arias Pérez, Karim-Jean Armache, Fréderic Chédin, Nicole J. Francis

## Abstract

Epigenetic regulation is conveyed through information encoded by specific chromatin features. Non-canonical nucleic acid structures could in principle also convey biological information but their role(s) in epigenetic regulation is not known. Polycomb Group (PcG) proteins form memory of transient transcriptional repression events that is necessary for development. In *Drosophila*, PcG proteins are recruited to specific DNA sequences, Polycomb Response Elements (PREs). PREs are switchable memory elements that can exist in repressed, active, or unengaged states ^1,2^. How PcG activities are targeted to PREs to maintain repressed states only in appropriate developmental contexts has been difficult to elucidate. Biochemically, PcG protein complexes modify chromatin to maintain gene repression ^1,3,4^. However, PcG proteins also interact with both RNA and DNA, and RNA is implicated in the targeting of PcG function. We find that R-loops, three-stranded nucleic acid structures formed when an RNA hybridizes to its complementary DNA and displaces the other DNA strand ^5^, form at many PREs in *Drosophila* embryos, and correlate with the repressive state. R-loops are recognized by the PcG complex PRC1 in vitro. Unexpectedly, we find that the PcG complex PRC2 has RNA strand invasion activity, which can drive formation of RNA-DNA hybrids, the key component of R-loops. Our results suggest a new mechanism for targeting PcG function through R-loop formation by PRC2 and recognition by PRC1. More generally, our findings suggest formation and recognition ^6^ of non-canonical nucleic acid structures as an epigenetic mechanism.

## Main Text

During *Drosophila* embryogenesis, transiently expressed transcription factors activate homeotic (*Hox*) genes in certain regions of the embryo and repress them in others to dictate the future body plan ^7^. Polycomb Group (PcG) proteins form a memory of these early cues by maintaining patterns of *Hox* gene repression for the rest of development ^1,7,8^. This paradigm for transcriptional memory is believed to be used by the PcG at many genes in *Drosophila*, and to underlie the conserved and essential functions of PcG proteins in cell differentiation and development from plants to mammals ^9,10^. Polycomb Response Elements (PREs) are DNA elements that can recruit PcG proteins, but they also recapitulate the memory function of the PcG— when combined with early acting, region-specific enhancers in transgenes, they maintain transgene repression in a PcG-dependent manner only in regions where the early enhancer was not active ^1,11,12^. PREs contain a high density of binding sites for transcription factors that can recruit PcG proteins through physical interactions ^12^. However, the widespread expression, binding pattern, and properties of factors that bind PREs cannot explain how PREs can exist in alternate, transcription-history dependent states to maintain restricted patterns of gene expression, or how they can switch between states ^1^. Furthermore, DNA sequences with PRE-like properties have been difficult to identify in other species ^12–14^ despite the conservation of PcG complexes, their biochemical activities, and their critical roles in development.

RNAs may provide context specificity to PcG protein recruitment and function. Some PREs, and some PcG binding sites in mammalian and plant cells, are transcribed into ncRNA, while others reside in gene bodies, and thus are transcribed when the gene is expressed ^15,16^. Both the direction and level of transcription have been correlated with the functional state of PREs ^15–17^. The PcG complex <u>P</u>olycomb <u>R</u>epressive <u>C</u>omplex 2 (PRC2) has a well-described high affinity for RNA ^18–22^. RNA is suggested to recruit PRC2 to specific chromatin sites ^18^, but RNA binding can also compete for chromatin binding and inhibit PRC2 activity ^16,22-24^. One way for RNA to interact with the genome is by the formation of R-loops, three-stranded nucleic acid structures formed when an RNA hybridizes to a complementary DNA strand, thereby displacing the second DNA strand ^5^. The formation of R-loops over genes with low to moderate expression is associated with increased PcG binding and H3K27 trimethylation (H3K27me3) in human cells ^25^ and R-loops have been implicated in promoting PcG recruitment in mammalian cells ^6^, although other evidence suggests they antagonize recruitment ^26^. We hypothesized that R-loop formation could biochemically link RNA to silencing through PREs and tested this idea in the *Drosophila* system.

To determine whether R-loops form at PREs, we carried out two biological replicates of strand-specific <u>D</u>NA-<u>R</u>NA Immuno<u>p</u>recipitation followed by next generation sequencing (DRIP-seq) in *Drosophila* embryos (2-6 and 10-14 hour (H)) and in S2 cells (Fig. 1, Extended Data Fig. 1). DRIP-seq peaks called relative to both input and RNaseH-treated control samples and present in both replicates were analyzed. 10 positive sites were validated by DRIP-qPCR (Extended Data Fig. 1b). About two thirds of R-loops form over gene bodies (Extended Data Fig. 1). R-loops are observed over genes encompassing all levels of transcription, although a majority are associated with genes with no or low levels of expression (Extended Data Fig. 2a, b). Most R-loops form with the strandedness expected from annotated transcripts (Fig. 1 a, c, Extended Data Fig. 2c), as observed in other species ^25,27,28^.

**Fig. 1.**
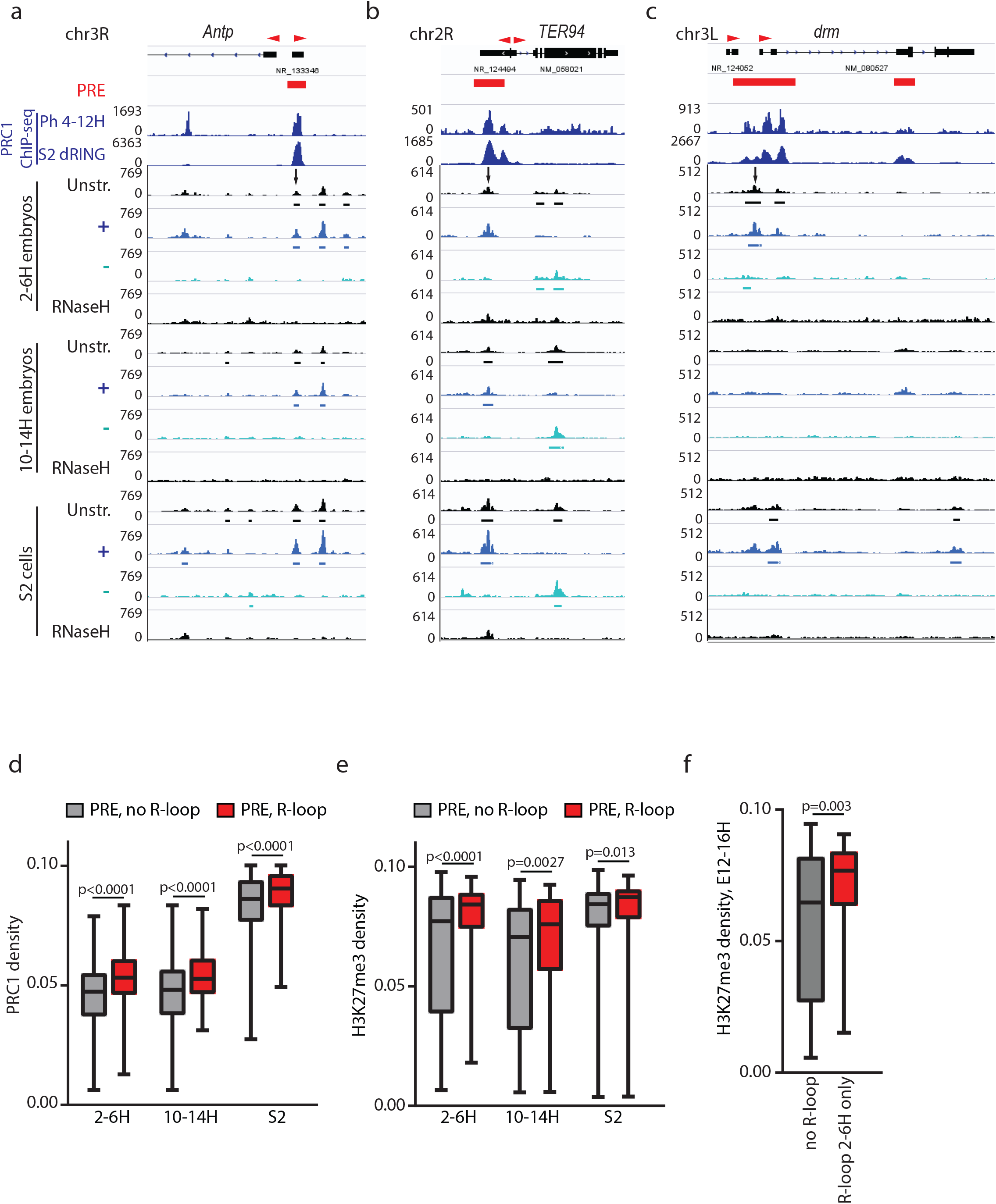
R-loops form at *Drosophila* PREs and correlate with a repressed state. a-c. DRIP-seq traces showing R-loop formation at PREs bound by PRC1 components (arrows) in 2-6H and 10-14H *Drosophila* embryos (Ph), and in S2 cells (dRING). RNaseH-treated samples are negative controls. “Unstr” indicates all R-loops, while + and – indicate strand specific tracks; direction refers to the DNA in the RNA-DNA hybrid. Called peaks are indicated under the traces. Red arrowheads above genes indicate direction of annotated transcripts. d, e. Median normalized intensity of PRC1 components (d), or H3K27me3 (e) over PREs with or without R-loops. 2-6H and 10-14H R-loop data are compared with Ph at 4-12H and H3K27me3 at 4-8H and 12-16H respectively. S2 cell R-loop data are compared with dRING. Whiskers show min. to max. f. Median normalized intensity of H3K27me3 at 12-16H over PREs where R-loop formation is detected in 2-6H but not in 10-14H compared to PREs with no R-loops detected at either stage. See also Extended Data Fig. 2.

We detect R-loops at 22-33% of PREs (Fig. 1 a-c, Extended Data Fig. 1d, 2 a-d). R-loops at PREs are more likely to have an antisense orientation to annotated transcripts, or have no overlapping annotated transcript, than total R-loops (Extended Data Fig. 2c). To test whether R-loops are related to the functional state of PREs, we compared PcG protein binding at R-loops that do or do not form R-loops in each of our three samples. For each PcG protein tested, the median read density over PREs with R-loops is higher than that for PREs without R-loops (Fig. 1d-f, Extended Data Fig. 2e-g). Although binding of PcG proteins to PREs is necessary for their repressive function, it may not be sufficient, since analyses of PcG protein binding at a small number of PREs in the ON and OFF states did not detect differences in PRC1 or PRC2 binding ^29,30^. Instead, histone modifications at and around PREs are correlated with the functional state so that PREs in the OFF state are marked with H3K27me3 ^29^. In both developing embryos and S2 cells, H3K27me3 density is higher at PREs with R-loops than those without R-loops (Fig. 1e). H3K27Ac, a mark of the active state, is found at a small number of PREs, but correlates weakly with the presence of R-loops (Extended data Fig. 2 d, h-j, 3d-f). To test whether transient presence of an R-loop at PREs predicts the repressed state, we analyzed PREs that form R-loops 2-6H embryos that are no longer detected in 10-14H embryos for the presence of H3k27me3 in later embryonic states (12-16H). PREs that formed R-loops in early embryos have a higher density of H3K27me3 at subsequent developmental stages than PREs that do not form R-loops at either stage (Fig. 1f), and are not enriched for H3K27Ac (p=0.0885).

To understand biochemically how R-loops could promote the repressive state of PREs, we turned to in vitro assays. We prepared R-loops *in vitro* by transcribing templates containing a PRE sequence (Fig. 2a). R-loops are visualized as bands containing radiolabelled RNA that co-migrates with the DNA template, and their identity confirmed by their sensitivity to RNase H and DNase I, and resistance to RNaseA (Fig. 2b, Extended Data Fig. 4a, b). We incubated either of the two main PcG complexes, PRC1 or PRC2, (Extended Data Fig. 4c,d) with transcribed templates, and fractionated the reactions by sucrose gradient sedimentation. Nucleic acids bound by PcG complexes sediment near the bottom of the gradient and unbound nucleic near the top (Fig. 2a). PRC1 binds preferentially to R-loop containing templates while PRC2 shows no preference (Fig. 2c-f, Extended Data Fig. 4e-g). Both complexes bind tightly to RNA (Extended Data Fig. 5a-e).

**Fig. 2.**
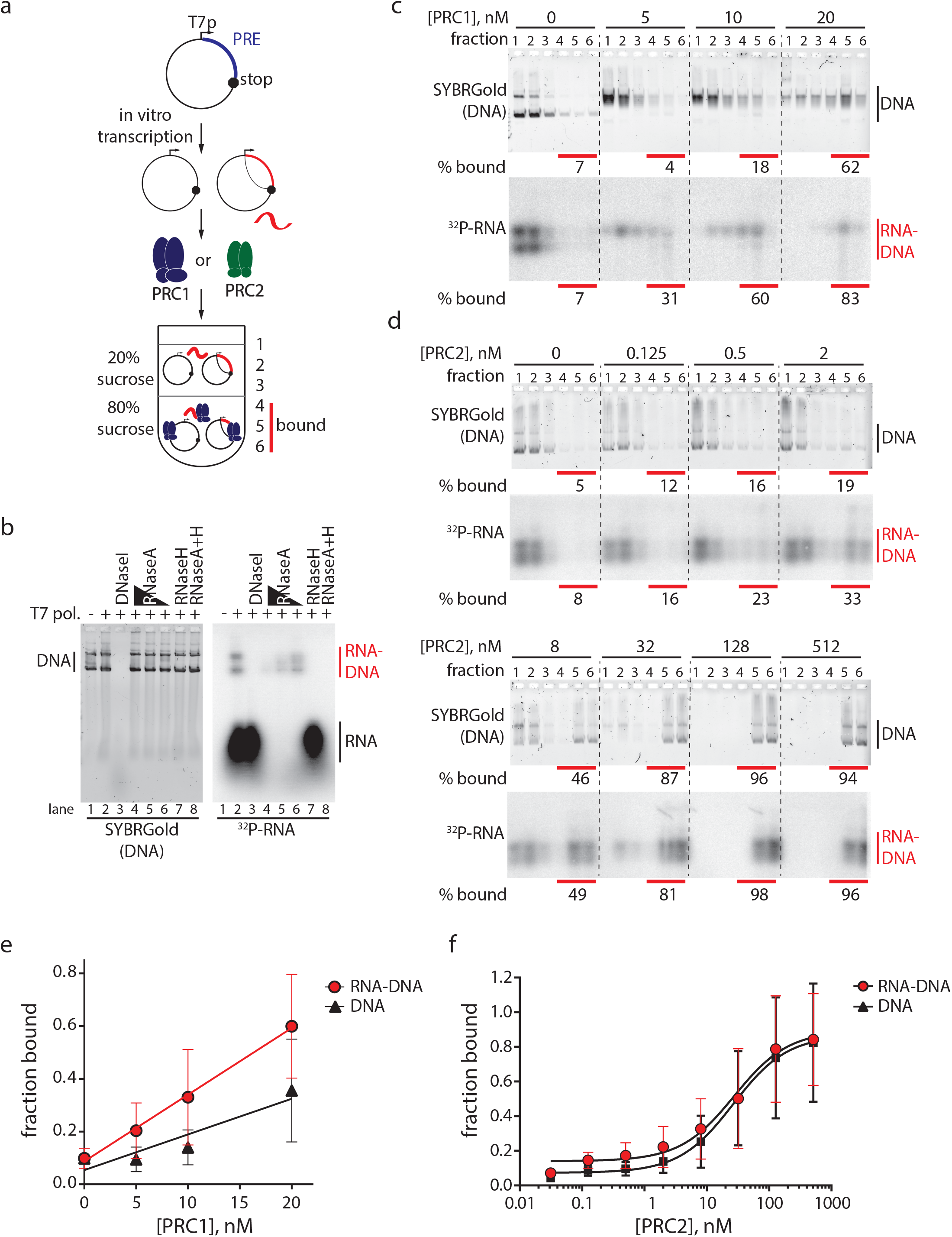
PcG complexes recognize R-loop containing templates. a. Scheme for testing binding of PcG complexes to R-loop containing plasmids. b. Radiolabelled RNAs that co-migrate with the DNA plasmid (lane 2) are confirmed as R-loops by their sensitivity to DNaseI (lane3) and RNaseH (lane 4), and resistance to RNaseA (lanes 5-7). c, d. Representative gradients of PRC1 (c) and PRC2 (d) binding. ^32^P-RNA panel shows RNaseA digested RNA-DNA hybrids. See Extended Data Fig. 4 for full gels. E, F. Quantification for PRC1 (e, n=8) and PRC2 (f, n=6). Points show the mean +/− S.D. RNA-DNA and DNA curve fits are different for PRC1 (p<0.0001, exact sum-of-squares F-test).

A small increase in R-loops is observed in some experiments in which PRC2 is incubated with transcribed templates, suggesting PRC2 might influence R-loop formation (Extended Data Fig. 5f,g). To test this, we mixed purified radio- or fluorescently-labelled RNA with dsDNA templates and titrated in PRC2 (Fig. 3a-c). We observe PRC2 dose-dependent appearance of a labelled RNA that migrates at the position of dsDNA (Fig. 3d, e, g, Extended Data Fig. 6a, b). These putative strand invasion products form with either the sense or anti-sense RNA, but not with a non-complementary RNA, indicating that base pairing between RNA and DNA is required (Fig. 3d-f). The products are sensitive to RNaseH and resistant to RNaseA, confirming that they contain RNA-DNA hybrids (Fig. 3h, i, Extended Data Fig. 6b). Two control proteins, the transcription factor NFY and the PcG protein Sxc, do not form strand invasion products, although they bind both DNA and RNA (Extended Data Fig. 6d-h). By the end of a 60-minute reaction containing 3 fmol of linear DNA and 1.9 fmol of RNA, as much as 60% of the RNA is incorporated into RNA-DNA hybrids, corresponding to 38% of the DNA molecules (Fig. 4a, b). Strand invasion activity of PRC2 does not require addition of nucleotides, but does require MgCl_2_ (Extended Data Fig. S6c). Finally, PRC2 strand invasion activity co-fractionates with PRC2 through size exclusion chromatography (Extended Data Fig. 7a-c). We conclude that PRC2 induces RNA strand invasion of dsDNA to produce RNA-DNA hybrid containing structures, the key component of R-loops.

**Fig. 3.**
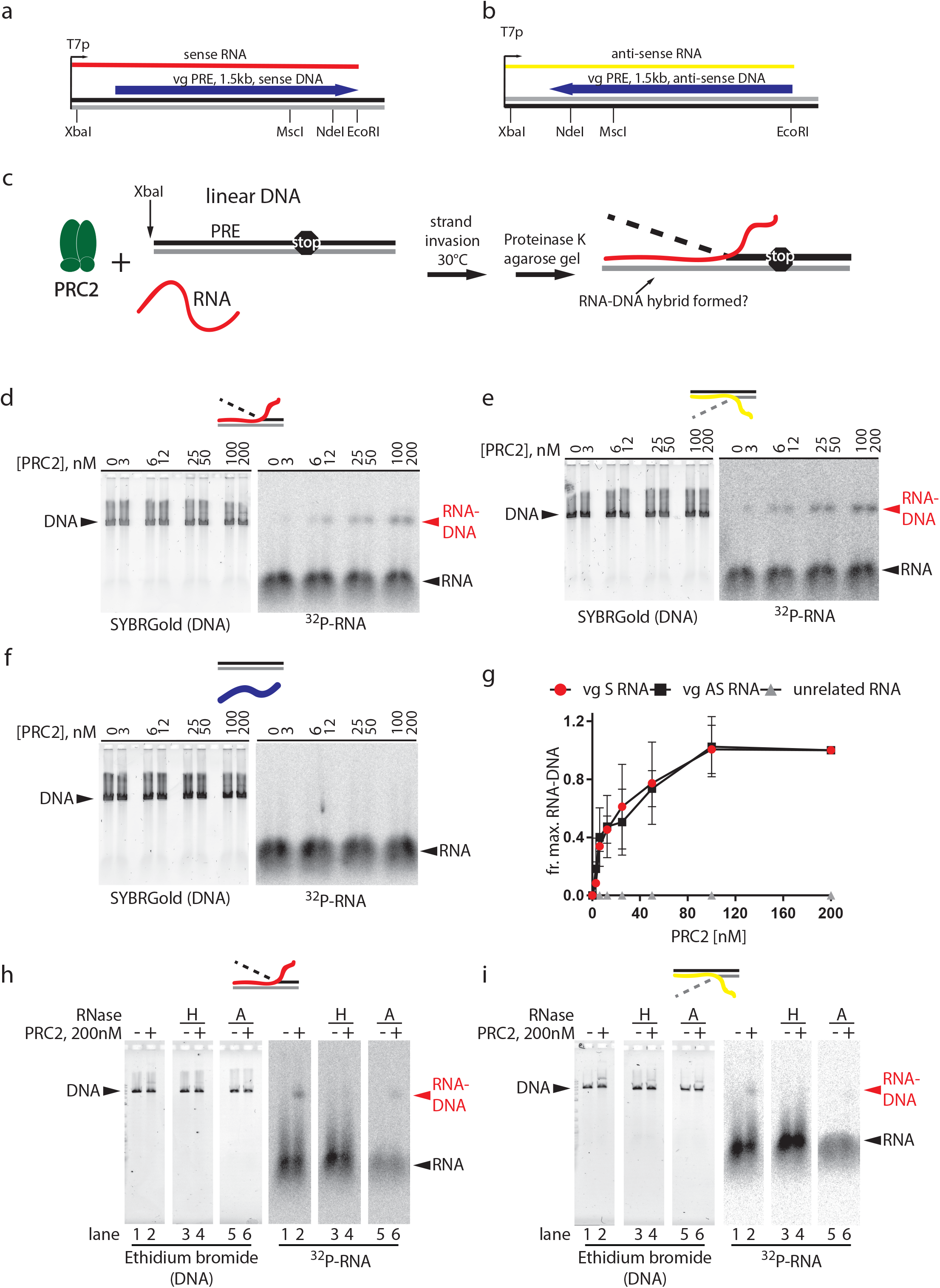
PRC2 has RNA strand invasion activity. a, b. Nucleic acids used for RNA strand invasion. Plasmids with the *vestigial* (*vg*) PRE in either orientation are transcribed to produce sense (a) or anti-sense RNAs (b). Linearized plasmid is used as the DNA template. c. Strand invasion assay scheme. d-f. Titration of PRC2 with either the sense (d) or anti-sense (e) *vg* PRE RNA, or a non-complementary RNA (f). g. Quantification of 3 titrations; graphs show mean +/− S.D of the the fraction of signal with 200 nM PRC2. h, i. RNA strand invasion products are sensitive to RNaseH (lane 4) but resistant to RNaseA (lane 6). See also Extended Data Fig. 6.

**Fig. 4.**
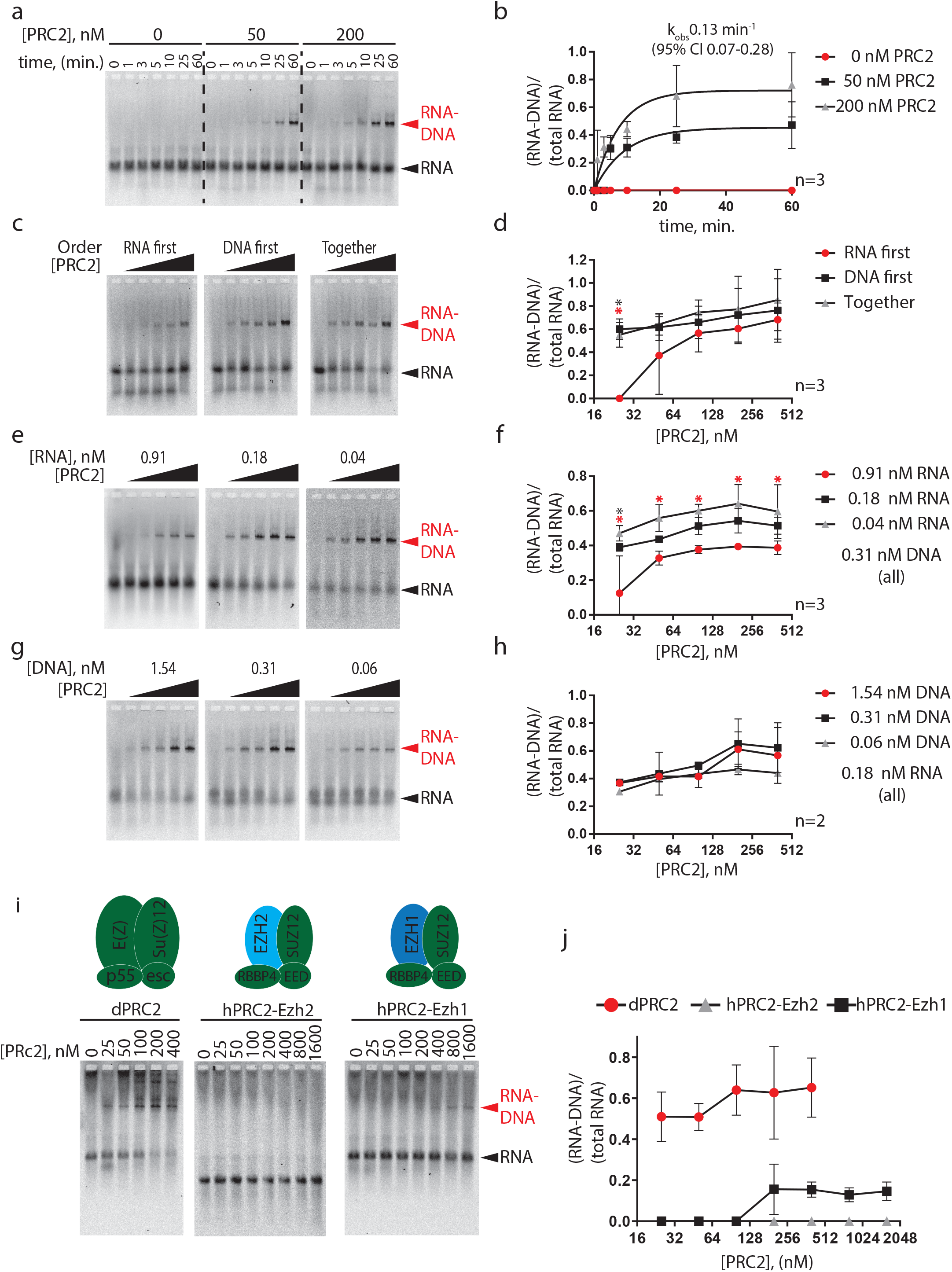
Characteristics and conservation of PRC2 strand invasion activity. In a-h, left panel shows a representative gel of Cy5-labelled RNA, and graph (right panel) summarizes multiple experiments. a, b. Time course of RNA strand invasion. c, d. Effect of order of addition of nucleic acids. Asterisks in all panels indicate p<0.05 (unpaired student’s t-test). red=RNA vs. DNA; black=RNA vs. together. c-g. Effect of increasing RNA (e, f) or DNA (g, h). Asterisks: red=0.91 nM vs. 0.04 nM; black=0.91 nM vs. 0.18 nM. I-All PRC2 titrations are 25-400 nM. i, j. PRC2-EZH1 has strand invasion activity. See also Extended Data Fig. 8.

PRC2-mediated strand invasion could require binding to DNA, to RNA, or to both. Detailed analyses of PRC2 binding to nucleic acids and chromatin are consistent with PRC2 making multiple contacts with both substrates ^24,31^, while functional assays are consistent with a single binding site that can bind chromatin, DNA or RNA but has highest affinity for RNA ^16,22,23^. To understand the role of RNA and DNA interactions in PRC2-mediated strand invasion, we manipulated the reaction conditions. Addition of RNA prior to DNA, or increasing the amount of RNA, inhibits the reaction, while adding DNA prior to RNA or increasing DNA has little effect (Fig. 4c-h, Extended Data Fig. 8a-d). These data are consistent with PRC2-DNA interactions being critical for strand invasion, and higher affinity PRC2-RNA interactions competing for them. This proposed mechanism, in which the protein binds the dsDNA template, resembles “inverse RNA strand invasion” described for the repair proteins Rad52 and RecA ^32–34^ (See Extended Data Fig. 9a-c for models).

PRC2 and its methyltransferase activity are conserved from plants to human, as are connections between PRC2 and RNA ^9,16-18,35^. We therefore anticipated that strand invasion activity would be conserved. We find that PRC2-EZH1 has strand invasion activity at concentrations that coincide with its DNA binding activity, while PRC2-EZH2 has little activity under these conditions (Fig. 4i, j, Extended Data Fig. 8e).

The demonstration that PRC2 has RNA strand invasion activity, that PRC1 can recognize R-loops, and that R-loops are present at PREs in vivo suggest a mechanistic model for how RNAs can promote the off state of PREs through PRC2-driven R-loop formation. R-loops could synergize with PRE-binding proteins to recruit PRC1 and would also sequester the RNA and tether it to the genome, preventing it from competing with the chromatin substrate for PRC2 binding. Stabilization of PcG proteins at PREs through R-loop formation would promote chromatin modification through the well-known activities of PRC1 and PRC2 ^1,4^. R-loops may also interfere with binding or function of proteins that promote the active state of PREs. Our data indicate that both coding and ncRNAs form R-loops. The regulation of these RNAs and therefore of R-loops could provide transcription history and developmental context specificity to PcG recruitment by transcription factors that constitutively recognize PREs. A conceptually similar model for how high levels of RNA production at PREs could promote the ON state and low levels the OFF state was proposed previously ^17^, but R-loop formation provides a mechanism by which it can occur.

The connection between RNA and PRC2 has been recognized for some time, in species from plants to humans ^16-18,35^, but mechanisms beyond RNA binding by PRC2 have not previously been described. Our discovery of PRC2-mediated RNA strand invasion, and R-loop formation at PREs, suggests a mechanism to connect RNA to PcG targeting and function, and the formation and recognition ^6^ of non-canonical nucleic acid structures to epigenetics (Extended Data 10).

## Acknowledgements

We thank E. Lécuyer’s lab for assistance collecting *Drosophila* embryos, O. Neyret for advice on preparation of NGS samples, J. Mallette for technical assistance, C. Gentile for advice on bioinformatics, M. Wilson for S9.6 antibody, M. Reijns for plasmid to express hRNaseH2, M. Drolet for intellectual support, and F. Robert, M. Drolet, and members of the Francis lab for comments on the manuscript. This research was enabled in part by support provided by Calcul Québec (www.calculquebec.ca) and Compute Canada (www.computecanada.ca).

## Funding

Work in the N.J.F. lab was funded by CIHR 311557, in the K.J.A. lab by a grant from the David and Lucile Packard Foundation, NIH 5R01GM115882-03, and 5T32HL007151-40 (to D.G.), and in the F.C. lab by NIH R01-GM120607.

## Author contributions

Conceptualization: N.J.F. & C.A.; Investigation: C.A., O.A. & N.J.F.; Formal analysis: C.A. & N.J.F.; Resources: V.C. & D.G., K.J.A., L.A.S., F.C.; Writing: N.J.F., C.A., K.J.A., F. C., Supervision: N.J.F., Funding acquisition: N.J.F., F.C., K.J.A.

## Competing interests

The authors declare no competing interests.

## Materials & Correspondence

All correspondence and material requests should be directed to N.J.F.

