## Supplementary information for "RNA strand invasion activity of the Polycomb complex PRC2"

**This PDF file includes:**

Materials & Methods
References 36-58
Extended Data Fig. 1-10

**Materials and Methods:**

**Protein expression and purification:**

Human RNaseH1: A 6X-His tag was added to MBP-hRNaseH1, which was expressed in and purified from *E. coli* based on a previously described protocol [39,40], except that Ni-NTA beads were used for the first step instead of amylose beads.

hRNaseH2: The RNaseH2 trimer was produced using the multi-cistronic pMAR22 vector essentially as described [40].

PRC1, PRC2, dSxc, hNFY: PRC1 and PRC2 were expressed in and purified from Sf9 cells, with the following modifications to previously published protocols for anti-FLAG affinity purification [41-43]. For PRC1, nuclear extracts were prepared from Sf9 cells infected with viruses for the 4 subunits [43] but nuclei were purified through a sucrose cushion prior to nuclear extraction. During the purification, the 2 M KCl wash in the published protocol was replaced with a wash consisting of BC2000N + 1 M Urea (20 mM Hepes, pH 7.9, 2 0.4 mM EDTA, 2 M KCl, 1 M deionized urea, 0.05% NP40, no glycerol). Additionally, prior to eluting the protein, anti-FLAG beads were incubated 3-5 volumes of BC300N with 4 mM ATP + 4 mM $MgCl_2$ for 30 min. at room temperature. This step reduces the amount of HSC-70 that co-purifies with PRC1.

For PRC2 expression and purification, E(Z) was tagged with 6-His, and either Esc or Su(Z)12 with FLAG, and bacculovirus infected Sf9 cells were harvested after 3 days. PRC2 was purified by anti-FLAG affinity as described [42] followed by Ni-NTA. FLAG peptide elutions were carried out in BC300 without EDTA or DTT. FLAG elutions were passed over Ni-NTA beads twice, beads were washed with 30 volumes of BC300 (without EDTA or DTT) and eluted in BC300 + 250 mM Imidazole. Eluted protein was pooled and dialyzed through 3 changes of

BC300 with EDTA, PMSF, and DTT. PRC2 was concentrated to ~1 mg/ml, NP40 was added to 0.05%, and protein was stored at -80$^0$ C.

Extract preparation and anti-Flag purification of F-Sxc and F-NFY were as described for PRC2. hPRC2-EZH1 and hPR2-EZH2 were prepared as described [44].

**Plasmids for R-loop formation and strand invasion:** PRE sequences were amplified by PCR from *Drosophila* genomic DNA and cloned into the pET-Blue1 vector (Millipore) downstream of the T7 promoter. Detailed maps are available on request. For strand invasion, plasmids were digested with a single restriction enzyme, purified by phenol-chloroform extraction, and ethanol precipitated.

**R-loop formation in vitro:** In vitro transcription was carried out in 50 µL reactions with 300 ng of DNA in 40 mM Tris-HCl pH 8.0, 8 mM $MgCl_2$, 25 mM NaCl, 2 mM spermidine, 30 mM DTT, 40 nM ATP, 40 nM CTP, 40 nM GTP, 8 nM UTP (New England Biolabs, NEB), 2.6 nmol of radiolabelled UTP (PerkinElmer) and 6 U of T7 RNA polymerase (NEB) for 30 min at 30$^0$ C. Polymerase was heat inactivated for 10 min. at 65$^0$ C. Each sample was split and incubated with 2 U of DNaseI in DNaseI buffer (NEB), 5 U of RNaseH in RNaseH buffer (NEB), 1 ng, 100 pg or 10 pg of RNaseA (Qiagen) or 5 U of RNAseH and 100 pg of RNaseA for 1 H at 30$^0$ C. After digestion with 3 µL of DSB-PK (6.7 µg/µL of proteinase K (Biobasic), 1% SDS, 50 mM Tris-HCl pH 8.0, 25% glycerol and 100mM EDTA) for 1H at 50$^0$ C, nucleic acids were resolved on 1% agarose .5X TBE gels and stained with SYBRGold (Thermo Fisher) followed by transfer to HYBOND membranes (GE Healthcare) and exposure to a phosphor imager screen.

In Extended Data Fig. 4a, b, in vitro transcription (1) was performed as described above with the omission of radiolabelled UTP. In vitro transcription (2) was performed with 300 ng of plasmid in a 50 µL reaction volume in presence of 20 mM DTT, 0.05 % Tween-20, 0.25 mM rNTP and 100 U of T3 or T7 RNA polymerase for 30 min at $37^0$ C followed by 10 min at $65^0$ C. 60 ng of transcribed template was treated with 10 U of RNaseH in presence of RNaseH buffer or with 1 ng of RNaseA supplemented with NaCl to 200 mM for 30 min at $37^0$ C. Reactions were stopped by addition of 20 µg of proteinase K and incubated 30 min at $37^0$ C. Samples were resolved on 1% agarose 0.5X TBE gel, stained with ethidium bromide and imaged on Typhoon Imager (GE Healthcare).

**Sucrose gradient sedimentation analysis of PcG complex binding to transcribed templates:** A 5 µl transcription reaction containing 25 ng of DNA template was carried out as described above. PRC1 or PRC2 were added and the reaction brought to a final composition of 180 mM KCl, 5 mM $MgCl_2$, 1 mM DTT and 50 ng/µl BSA (NEB) in a 20 µL final volume. Reactions were incubated 1 hour at $30^0$ C. The whole sample was then loaded onto a step gradient composed of 50 µL of 80% sucrose in BC150 and 150 µL of 20% sucrose in BC150 and. Gradients were centrifuged 50 min. at 40,000 rpm in a TLS 55 rotor (Beckman) and resolved into 6 fractions. One half of each fraction was treated with 5 µL of DSB-PK overnight, and the other half was treated with 2 ng of RNaseA in presence of 330 mM NaCl for 2 hours at $30^0$ C before proteinase K digestion. Samples were then resolved on 1% or 0.8% (RNaseA treated samples) agarose 0.5X TBE gels, stained with SYBRGold, transferred to a HYBOND membrane, and exposed to a phosphor imager screen.

**RNA production and labelling:** RNAs were produced from linear templates using the Ampliscribe T7-flash transcription kit (Lucigen) using the manufacturer’s protocol in the presence of 25 mM of amino-allyl UTP (Sigma). After purification, RNAs were labelled with NHS-Cyanin-5 (Kerafast) in 70 mM $NaHCO_3$ pH 8.8 with murine RNase inhibitor (NEB) for 2 hours at RT. RNAs were then precipitated with 0.3 M sodium acetate pH 5.3, glycogen and ethanol, washed with 70% ethanol, and resuspended in TE. RNAs were passed through a G50 column equilibrated with TE. The quality of labelled RNA and efficient removal of free dye were determined by loading the RNA on agarose gels.

Radiolabelled RNAs were produced from circular templates by transcribing 600 ng of DNA in RNA polymerase buffer (NEB), 1 mM DTT, 625 µM of rNTP, 6.5 nmol of radiolabelled UTP, 200 U of RNase inhibitor, and 250 U of T7 or T3 RNA polymerases (NEB) in 100 µL O.N. at $37^0$ C. DNA was removed from the reaction by adding 4 U of DNaseI and incubating 2 hours at $37^0$ C. RNAs were extracted with phenol-chloroform, ethanol precipitated, washed with 70% ethanol, resuspend in TE, and stored at $-20^0$ C.

**Strand invasion:** PRC2, diluted in BC300N was incubated with the indicated amount of DNA and fluorescent- or radio- labelled RNA for 25 min. at $30^0$ C in 180 mM KCl, 5 mM $MgCl_2$, 1 mM DTT and with 50 ng/ µL BSA in 10 µL reaction. After incubation, samples were treated with 3 µL of DSB-PK for 30 min. at $50^0$ C and resolved on 0.8% agarose 0.5X TBE gel. Gels were stained with SYBRGold or ethidium bromide, and imaged on a Typhoon Imager. For experiments with radio-labelled RNA, gels were transferred to HYBOND membrane and exposed to a phosphor imager screen.

For nuclease treatment of strand invasion products without phenol-chloroform extraction, after incubation with PRC2, samples were treated immediately with nucleases. For RNaseH treatment, 10X RNaseH buffer was added to a final concentration of 1X, followed by 2.5 U (radio-labelled RNA) or 1.25 U (fluorescently labelled RNA) of RNaseH. For RNaseA treatment, reactions were supplemented with 500 mM NaCl and 50 pg of RNaseA were added. Reactions were incubated for 30 min. at $30^0$ C. For phenol-chloroform extracted samples, reactions were stopped with 3 μL of DSB-PK and incubated for 30 min. at $50^0$ C. Nucleic acids were extracted with phenol-chloroform followed by ethanol precipitation and resuspension in TE. Nuclease digestion was carried out as described above. Nuclease digestions were stopped by the addition of 3 μL of DSB-PK, and samples were incubated 30 min. at $50^0$ C before analyzing on agarose gels.

When the order of DNA and RNA addition was tested, the first nucleic acid was added to PRC2 for 10 min. at 30°C before the addition of the second.

To test the activity of human PRC2, human or *Drosophila* PRC2 were diluted in 20 mM Tris-HCl pH 7.5, 300 mM NaCl, 1 mM DTT and 10% glycerol, then incubated with the 12.3 fmol of DNA and 1.8 fmol of fluorescently labelled RNA for 25 min at $30^0$ C in 50 mM NaCl, 50 mM Tris-HCl pH 8.0, 5 mM MgCl2, 1 mM DTT and 50 ng BSA in a 10 µL reaction. After incubation, samples were treated with DSB-PK for 30 min. at $50^0$ C and resolved on 0.8% agarose 0.5X TBE gels. Gels were stained with SYBRGold and imaged on a Typhoon Imager.

**EMSA:** EMSA were performed using the same conditions as for strand invasion. For hPRC2, optimized conditions described above were used. Linear DNA was used at 3 nM, and RNA at 1.8 nM. After 25 min. incubation at $30^0$ C, samples were loaded on 0.8% agarose 0.5X TBE gel and run at $4^0$ C. Gels were stained with SYBRGold or ethidium bromide, and imaged on a Typhoon Imager.

**Gel quantification:** For quantification of DNA and RNA-DNA hybrids from phoshpor imager and SYBR gold scans using ImageQuant, RNaseA treated samples were used. In cases where gel flaws obscured quantification of a lane, the gradient was excluded from analysis. Background subtraction was done using the rolling ball method. For band selection, the smallest possible “fixed width” bands that capture the whole signal were set for each gradient. These bands were placed in each lane so that every fraction was quantified. The signal from the bottom three fractions was divided by that for the total of the gradient for the fraction bound.

Strand invasion gels of Cy5 labelled RNA were imaged using Typhoon Imager (GE Healthcare) were quantified using ImageQuant (GE Healthcare). Lanes were created manually, then background was removed using minimum profile method and bands were identified manually.

**S2 cell culture:** *Drosophila* S2 cells were purchased from Invitrogen, and grown in Schneider’s media (Invitrogen) with 10% heat inactivated, insect cell tested FBS (Invitrogen). Cells were cultured at $27^0$ C in suspension in shaking flasks.

***Drosophila* collection:** Oregon R flies were grown at $25^0$ C. Embryos were collected on apple juice places and dechorionated for 2 min. in 50% bleach before being washed with $H_2O$ and stored at $-80^0$ C.

**Total nucleic acid extraction from S2 cells**: $8*10^7$ S2 cells were washed with 1X PBS and resuspended in 10 mL TE. Cells were lysed O.N. at $37^0$ C in presence of 0.5% SDS and 62.5 µg/mL of proteinase K. After phenol-chloroform-isoamyl acohol extraction, total nucleic acids were precipitated in the presence of 0.3 M sodium acetate pH 5.2 and 2.4 volume of 100% ethanol. Nucleic acids were then washed carefully 5 times with 70% ethanol, and resuspended in TE.

**Total nucleic acid extraction from *Drosophila* embryos:** Total nucleic acids were extracted from 500 uL of Oregon R embryos as described in Ejsmont et al., 2009 [45] with the omission of RNaseA. After precipitation the nucleic acids were washed carefully 5 times with 70% ethanol, and resuspended in TE. This material was subsequently processed for DRIP analysis as described below.

**DRIP-seq and DRIP-qPCR**: The DRIP protocol was adapted from Ginno et al. [36]. 500 µg of total nucleic acid were divided in 3 and each treated with 150 µg of RNaseA in presence of 0.5 M NaCl for 3 hours at $37^0$ C. gDNA was purified by phenol-chloroform extraction followed by ethanol precipitation and sonicated to 300 bp using a Covaris. Fragmented gDNA was treated with 2 U of RNaseIII [27] (Thermo Fisher) +/- 10 µg each of homemade RNaseH I and RNaseH II overnight at $37^0$ C. Immunoprecipitation was performed as described in Ginno et al. After elution, samples were purified with a PCR clean-up column (Macherey-Nagel) with NTB buffer to get rid of SDS followed by a DNA clean and concentrator column (Zymo Research). For sequencing library preparation, material from three immunoprecipitations were pooled. Libraries were prepared using the NEB next Ultra II kit for a directional library for Illumina (NEB). For strand specific DNA sequencing of the RNA-DNA hybrids, we started at the second strand synthesis step and ligated with NEB-next mutilplex oligos for Illumina (NEB). Paired-end sequencing was performed on an Illumina HiSeq 2500 at Genome Quebec.

For qPCR, input was diluted 10-fold and IPs 2-fold in water. PCR was carried out in 5 µl reactions consisting of 2 µl DNA, 2.5 µl PowerUp SYBR Green master mix (Thermo Fisher) and 0.25 µl of a 1 µM stock of each primer diluted in water. Standard curves were generated using a log titration of Drosophila genomic DNA purified from S2 cells (25 to 0.025 ng). Data were collected using a Viaa7 PCR system (Thermo Fisher) with 40 cycles. The standard curve was used to calculate DNA amounts. All standard curves had $R^2$ values of 0.9 or higher. Oligonucleotides used for qPCR [29,46] are list listed in Supplementary Table 1.

**DRIP-seq analysis**: FastQ files of DRIP-seq reads were trimmed with Trimmomatic (PE – phred33), using the GenPpipes ChIP-seq pipeline (steps 1-3) [47]. Reads with both mate pairs were aligned to the dm3 version of the *Drosophila* genome using Bowtie2/2.3.1(--fr --no-mixed --no-unal) [48]. Sam files generated by Bowtie2 were converted to bam, sorted and indexed (samtools (v. 1.4.1) [49] and Picard (http://broadinstitute.github.io/picard) MarkDuplicates (default parameters) was used to remove duplicates. To generate strand specific bam files, samtools was used as follows:

Forward strand: samtools view –f 99; samtools view –f 147, followed by samtools merge

Reverse strand: samtools view –f 83; samtools view –f 163, followed by samtools merge.

Peaks were called for DRIP versus input and DRIP versus RNaseH treated using MACS2[50] (v. 2.1.1) (-f BAMPE --bw 250 -g dm --mfold 10 30 -q 0.01). For strand specific peaks, strand specific files were used (e.g. F-strand DRIP, F-strand input, F-strand RNaseH). Peaks present in both DRIP vs. input and DRIP vs. RNaseH were retained (BEDTools intersect) [51] for each duplicate. Finally, BEDTools (intersect) was used to retain only peaks present in both duplicates, which were used for further analysis. The correlation between the replicates was examined using multiBigwigSummary on Galaxy (bin size: 1000 bp) followed by plotCorrelation using the Pearson correlation method. Correlations for replicates were: 2-6H 0.97, 10-14H 0.87, S2 0.99. Bigwig files were generated using DeepTools [52] v 2.5.3 (--binSize 10\ --normalizeUsingRPKM)

A list of PREs (Table S2) was generated by combining predicted PREs [53], PcG binding sites conserved through *Drosophila* species [54], and additional PREs from recent reports [16,46,55]. Multiple PREs predicted in the repeated histone gene clusters were removed, although ChiP-seq peaks for PcG proteins are observed at these sites. Finally, overlapping or touching PREs were merged (using BEDTools). The list of genomic coordinates for PREs is in Supplmentary Table 2.

To analyze overlaps between R-loops and PREs or other genomic elements, bed files of peak calls of unstranded, forward, and reverse strand peaks were merged to produce a consolidated set of R-loops. Overlap of R-loops or PREs with different genomic elements (Extended Data Fig. 1e-h) were generated with Pavis, with upstream and downstream regions both set at 5000 bp [56]. To correlate gene expression levels with R-loop formation (Extended Data Fig. 2a, b) the overlapping or closest gene to each R-loop was identified using BEDTools, ClosestBed on Galaxy. Level of gene expression were determined using RNA-seq data from embryos or S2 cells and genes were divided into categories based on their FPKM level (no to extremely low expression: FMPK<1, low expression: 1<FPKM<10, moderate expression: 10<FPKM<50 and high expression: FPKM>50). To compare R-loop orientation to annotated transripts, the “all EST” track was downloaded from UCSC, and BEDTools was used (intersect inervals, only overlaps occurring on the same strand).

To analyze overlap of PREs with PcG protein or H3K27me3 ChIP-seq peaks (Fig. 1d-f, Extended Data Fig. 2e-j), previously processed bed files were used with BEDTools (intersect). To analyze ChIP-seq signal intensity over PREs with and without R-loops, raw data (FASTQ files) were downloaded using the SRA tooolkit (v2.9.6) (http://ncbi.github.io/sra-tools/, SRA Toolkit Development Team), aligned with Bowtie2 as described above, duplicates removed (Picard), and RPKM-normalized bigwig files generated (DeepTools bamCoverage). BEDOPS[57] (v2.4.34) was used to convert bigwig files to wig and then bed files, and read densities quantified using BEDOPS bedmap (bedmap –count –echo-ref-name ). Read densities over each PRE were divided by the PRE length to obtain the final values. All data sets used to analyze R-loops are listed in Supplementary Table 3.

**Statistics & curve fitting:** Graphpad Prism was used for statistics and curve fitting. For time course data, the equation Y=ABmax*(1-exp[-k*X]) was used; for binding data, Y=ABmax*X/(X+Kd)+b was used. Fisher's exact test was used to compare curve fits. For students t-tests (Fig. 2), we used Holm-Sidak correction for multiple comparisons, alpha=0.05, with all points assumed to come from populations with the same S.D. For comparing distribution of ChIP-seq peaks or transcripts (Extended Data Fig. 2), data were organized as a contingency table (i.e. columns=peak/no peak, rows=R-loop/no R-loop) and compared using Fisher's exact test, reporting two-sided P-values. To compare ChIP-seq read intensities over PREs, Mann-Whitney tests were used, with 2-tailed p-values reported.

RegioneR [58] was used to conduct permutation tests of the overlaps between PREs and R-loops, or ChiP-seq peaks and PREs with and without R-loops (1000 permutations, randominze.function=randomizeRegions, evaluate.function=numOverlaps, count.once=TRUE, genome="dm3").

**Data Availability:** Sequence (DRIP-seq) data that support the findings of this study have been deposited in NCBI GEO with the accession code GSE127329. Other data that support the findings of this study are available from the corresponding author upon reasonable request.

**Extended Data Fig. 1 Overview of DRIP-seq analysis**. a. Distribution of DRIP-seq peak sizes. b. 10 positive sites and 3 negative sites were confirmed by DRP-qPCR using S2 cell genomic DNA with or without RNaseH treatment. c. Summary of overlaps of R-loops with genes. d. Summary of overlaps of R-loops with PREs. e-h. Pie charts of distribution of R-loops (e-g) and PREs (h) across the genome.

a

b

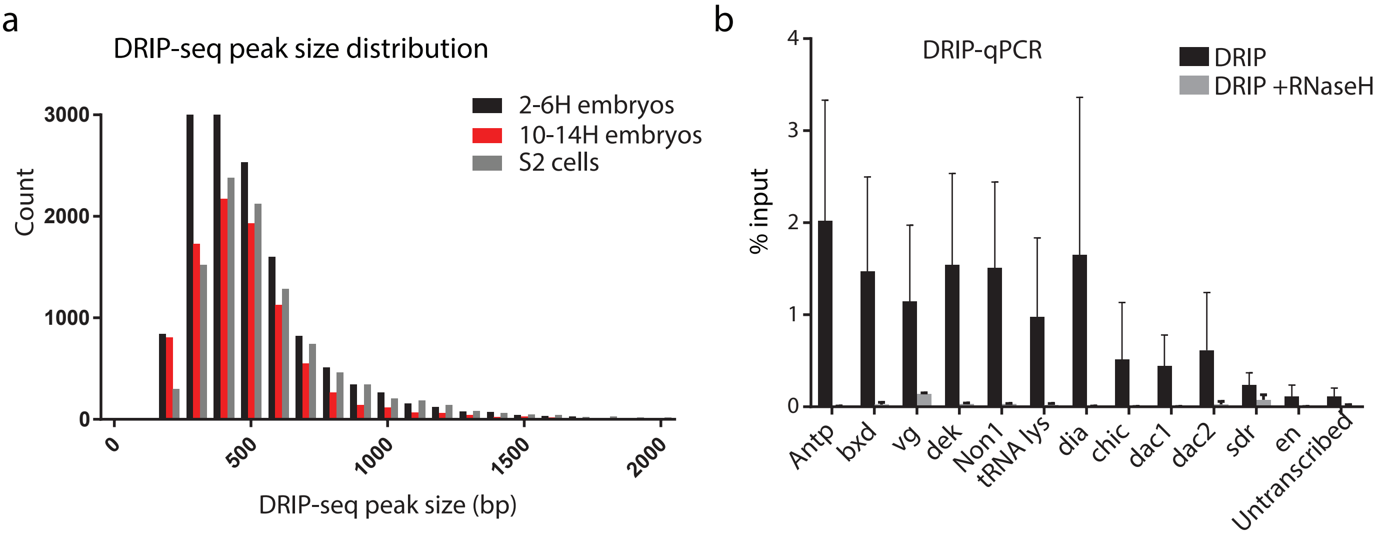


c

| | R-loop peaks | Overlap with gene body | Overlap with EST |
|---|---|---|---|
| 2-6H embryos | 17149 | 101460/17149 (61%) | 14709/17149 (86%) |
| 10-14H embryos | 10184 | 5779/10184 (57%) | 8497/10184 (83%) |
| S2 cells | 14549 | 9912/14549 (68%) | 11222/14549 (77%) |

d

| | PREs overlapping R-loops | p-value (permutation analysis) |
|---|---|---|
| 2-6H embryos | 180/541 (33%) | 0.001 |
| 10-14H embryos | 118/541 (22%) | 0.001 |
| S2 cells | 156/541 (29%) | 0.001 |

e

f

g

h

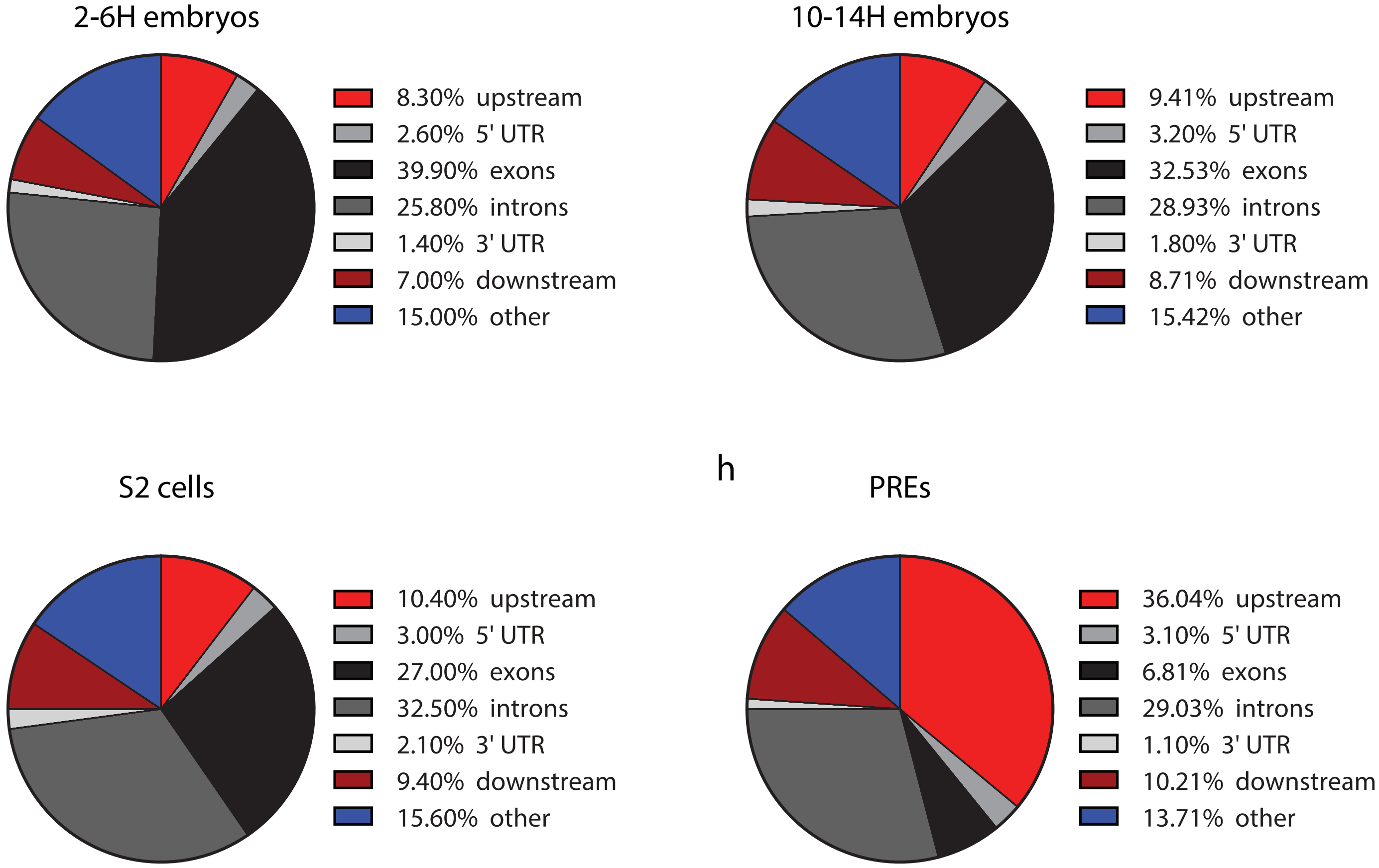


Alecki et al., Extended Data Fig. 1

**Extended Data Fig. 2 Relationship between R-loop formation, transcription, and PcG proteins.** a. Expression levels of all genes as compared with genes with R-loops. b. Expression level of genes associated with PREs that do not or do form R-loops. c. Strandedness of all R-loops or R-loops formed at PREs relative to annotated transcripts. p-values are for comparison of sense vs. antisense for R-loops that are transcribed. PREs also have a higher fraction of R-loops without annotated transcripts for all samples ($p<0.0001$ comparing transcribed versus untranscribed, Fisher's exact test). d. Summary of overlaps between PREs that do or do not form R-loops and H3K27me3 or H3K27Ac ChIP-seq peaks. Red numbers indicate that overlap is higher than random and blue numbers that it is lower than random. Overlap between PREs with or without R-loops is not significant, and is in fact anticorrelated in some cases (blue numbers). e-g. Density of normalized ChIP-seq reads over PREs that do or do not form R-loops. h-j. Fraction of PREs that do or do not form R-loops that overlap H3K27me3 or H3K27Ac peaks. p-values are for Fishers exact test comparing the overlap for PREs that do or do not have R-loops.

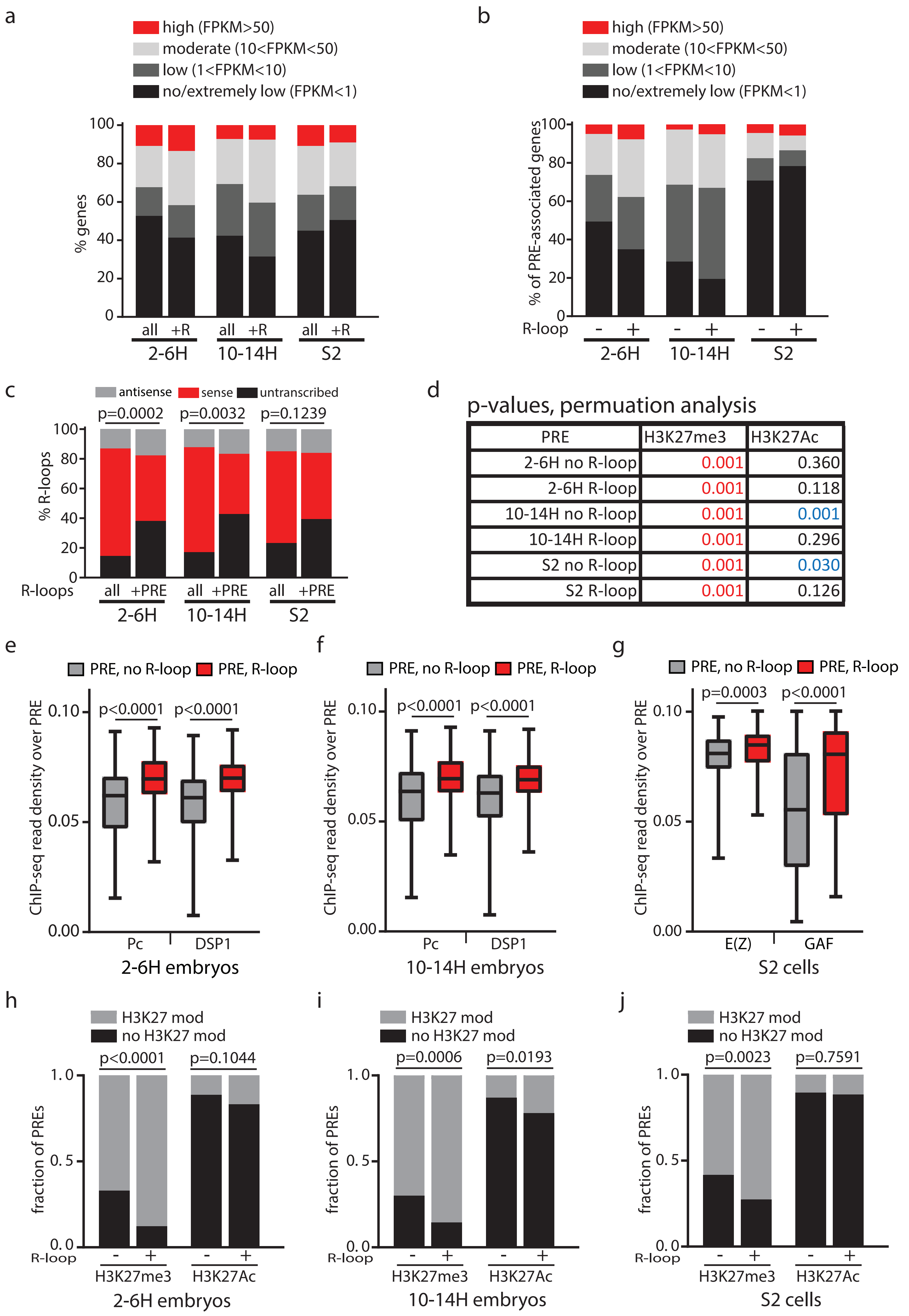


Alecki et al., Extended Data Fig. 2

**Extended Data Fig. 3 Additional examples of R-loops.** a. An R-loop forms over the PRC1-bound PRE with the same strandedness as *eve* transcription (arrow), but opposite to the direction of the overlapping ncRNA. b. An R-loop forms at a PRC1-bound PRE at the *Psc* gene. Note that an R-loop is detected at the PRE overlapping the promoter in 2-6H embryos (grey arrow), while R-loop signal is detected in all samples at the upstream PRE that overlaps the non-coding RNA (black arrow). c. Arrow indicates R-loop that overlaps a PRC1-bound PRE upstream of the *ben* gene but has no annotated transcript. d-f. Relationship between H3K27 modification, R-loops, and PREs in S2 cells. d. The *en* PRE has characteristics of the OFF state: the PRE has high levels of H3K27me3, both PRC1 and PRC2 binding, and forms R-loops (arrows). e. The PRE at the promoter for *CG12772* has characteristics of a balanced state, with both H3K27Ac and H3K27me3 modifications, PRC1 and PRC2 binding, and no R-loop (arrow). f. The PRE at *CG5953* has characteristics of the ON state, with PRC1 binding, high levels of H3K27Ac, no H3K27me3, and no R-loop formation. RNA-seq data (FPKM) are consistent with these states: *en*=0.016, *CG12772*=6.3, *CG5953*=3.6, 15.6 (2 isoforms).

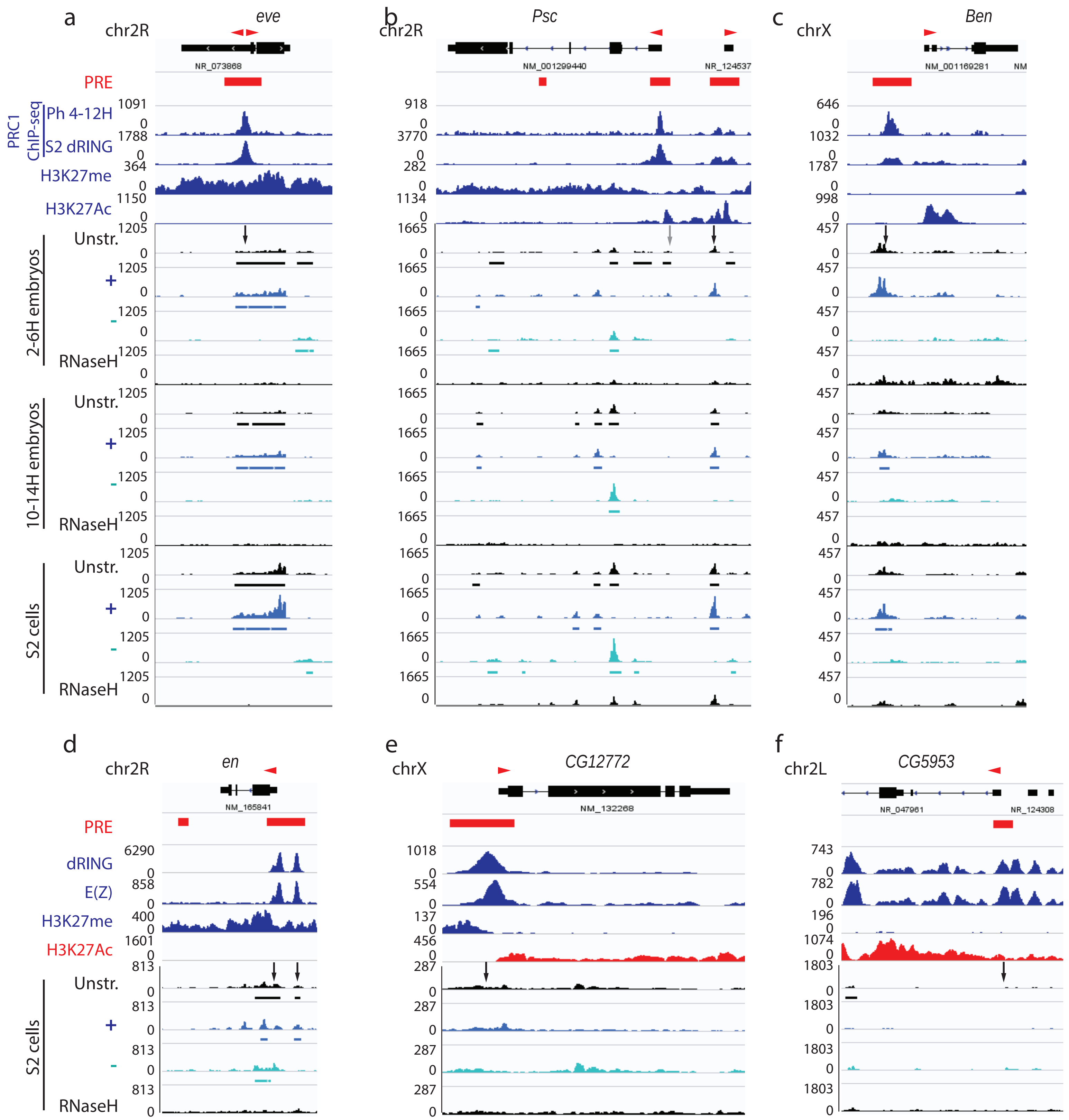


Alecki et al., Extended Data Fig. 3

**Extended Data Fig. 4 In vitro analysis of PcG-R-loop interactions.** a,b. Comparison of reaction conditions used to prepare R-loop containing templates for in vitro binding assays (set 1) and previously described conditions (2) [36]. a. *vg*-PRE containing plasmid template. b. pFC53, which contains a previously described strong R-loop forming human sequence that can be transcribed by T3 polymerase [36]. The templates we are using do not show the pronounced topological shift (compare lanes 1 and 2) observed with pFC53. This shift is dependent on R-loop formation since it is sensitive to RNaseH (lane 3). This could indicate that the *vg* PRE has less R-loop forming potential than pFC53. However, it could also indicate that the R-loops formed by the vg PRE are small (a few hundred bp) since the large size of the plasmid (~5kb) will obscure small topological shifts. Our conditions also result in much less transcription (compare RNA bands in conditions 1 and 2).R/N=relaxed/nicked; SC=supercoiled. Reactions were separated on agarose gels and stained with ethidium bromide. c,d. SYPRO Ruby-stained SDS-PAGE of PRC1 (c) (8% acrylamide) and PRC2 (d) (10% acrylamide) used in this study. PRC2 prepared with Flag-Su(Z)12 or Flag-Esc were used interchangeably. *=HSC70 contaminant. Molecular weight marker migration in kDa is indicated. e. Graph shown in Fig. 2e with all data points shown. Although the difference between binding to RNA-DNA hybrids and DNA is small, it was reproduced in all 8 experiments. f. Full gels of panels shown in Fig. 2c. g. Full gels of panels shown in Fig. 2d.

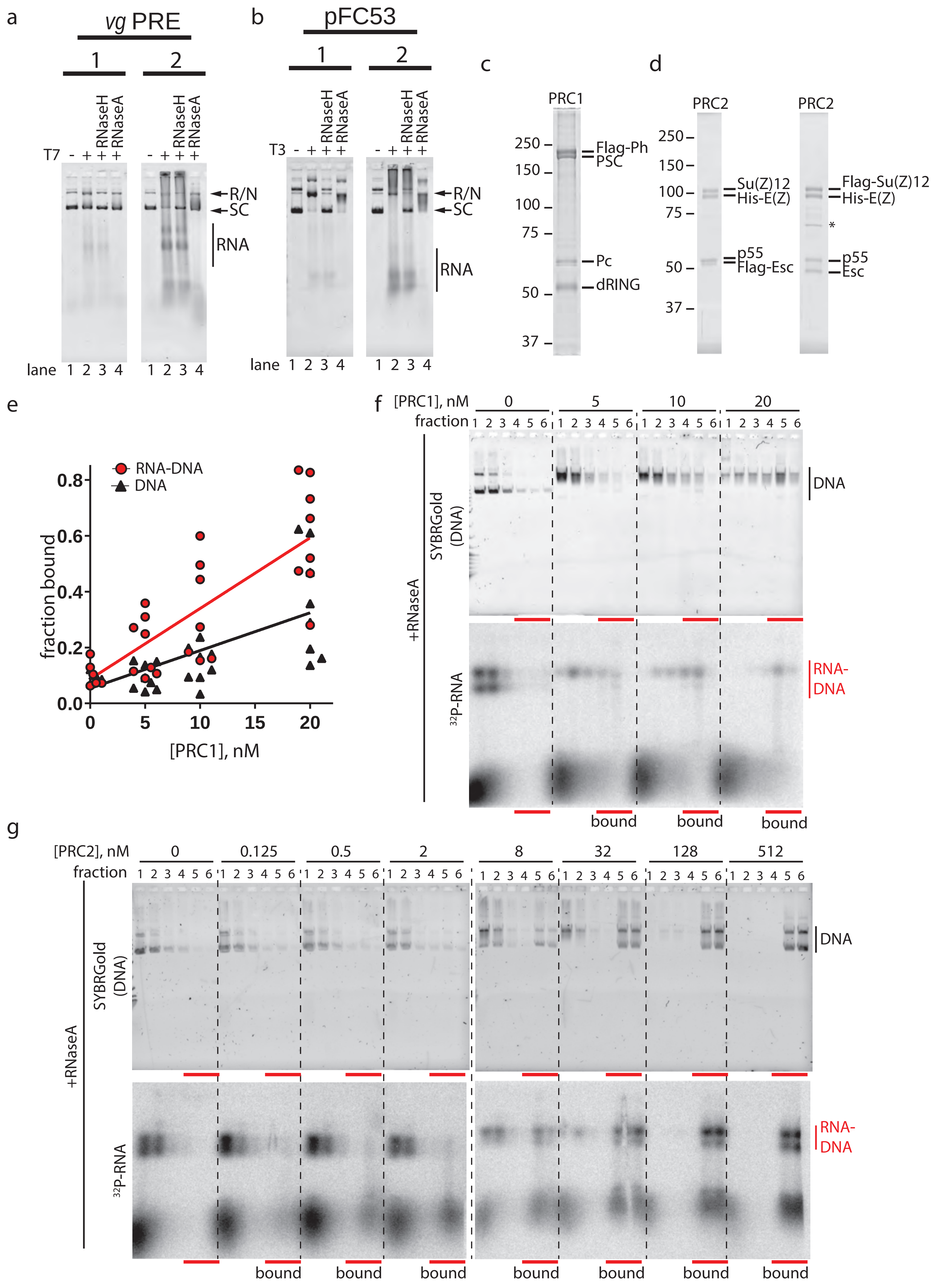


Alecki et al., Extended Data Fig. 4

**Extended Data Fig. 5 RNA binding by PRC1 and PRC2 and increased R-loops in the presence of PRC2.** a-c. Representative gels of EMSA for PRC1 (a) and PRC2 (b, c) binding to Cy5-labelledRNA (a, b) or unlabelled DNA (c) used in strand invasion assays. RNA was used at 1.8 nM, and DNA at 0.3nM. d, e. Summary of quantification of multiple EMSA experiments (PRC1 n=4, PRC2 n=6). Quantification was done by measuring the unbound (indicated by black arrowhead in a-c), and considering all signal above this band (indicated by the black line in a-c) as bound. Graphs show the mean +/- S.D. and fits are with non-linear regression. f. Schematic of assay to test the effect of PRC2 on R-loops formed by in vitro transcription of circular templates. After transcription by T7 polymerase, PRC2 is incubated with the transcription reaction. Reactions are digested with proteinase K and separated on agarose gels to assess R-loop levels. g. Gel of the effect of PRC2 on RNA-DNA hybrids showing a small increase in R-loop signal. We observed this activity in several experiments, with different preparations of PRC2, and it was the incentive for the strand invasion assay. However, the increase in RNA-DNA hybrids is generally less that 2-fold and many experiments do not show an increase. Using our strand invasion protocol, we have not yet been able to observe strand invasion with circular templates.

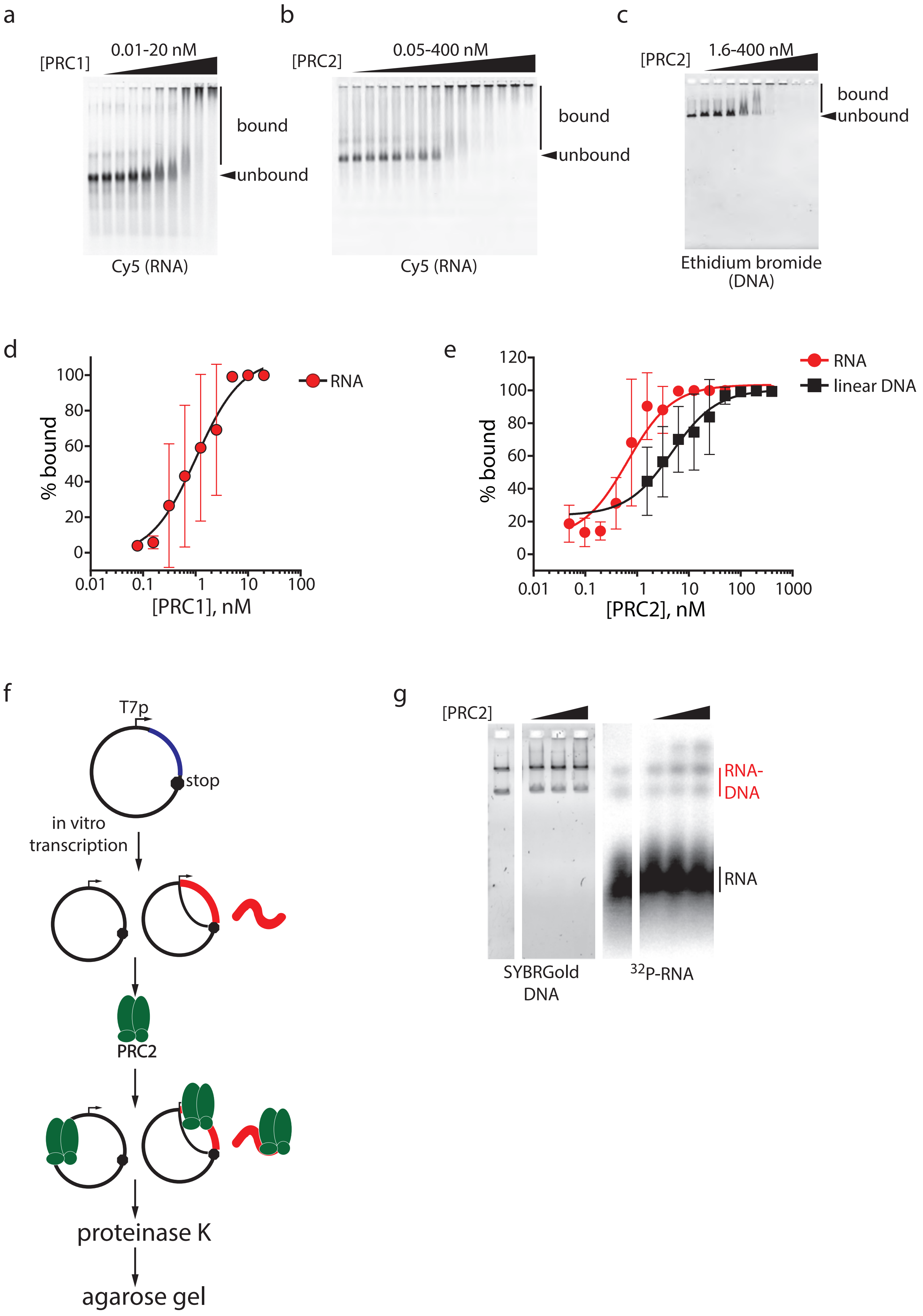


Alecki et al., Extended Data Fig. 5

**Extended Data Fig. 6 PRC2, but not other RNA and DNA binding proteins, mediates strand invasion with fluorescently labelled RNA**. a. Scheme for generating fluorescently labelled RNAs. DNA corresponding to the PRE sequence is amplified with primers, one of which contains the T7 promoter sequence for in vitro transcription. RNAs are produced by in vitro transcription in the presence of amino-allyl UTP, purified, and labelled with NHS-Cy5. These RNAs correspond precisely to the PRE sequence in the DNA template, and are used for strand invasion with linearized, PRE-containing plasmid DNA. Dashed lines indicate plasmid backbone sequence, which is not drawn to scale. Note that the radio-labelled RNAs used in Fig. 3 were generated by in vitro transcription of the circular plasmid template, so they all share ~200 bases of plasmid sequence between the T7 promoter and start of the PRE. b. Representative gels of PRC2 strand invasion activity with Cy5-labelled RNA. PRC2 strand invasion products (lanes 2, 3) are sensitive to RNaseH (lanes 4, 5). [PRC2]=50 and 200 nM, [DNA]=0.3 nM, and [RNA]=0.18nM. c. RNA strand invasion activity requires $MgCl_2$ but not ATP. Summary of 2 experiments. Error bars are S.D. d-h. Negative controls for strand invasion. d. SYPRO Ruby-stained SDS-PAGE gels of Flag-Sxc and Flag-NFY. *=HSC70 contaminant. e, f. EMSA demonstrating that both Sxc and NFY bind RNA (left) and linear DNA (right), the substrates for strand invasion. g, h. Neither Sxc nor NFY have strand invasion activity at concentrations where they can bind to both substrates.

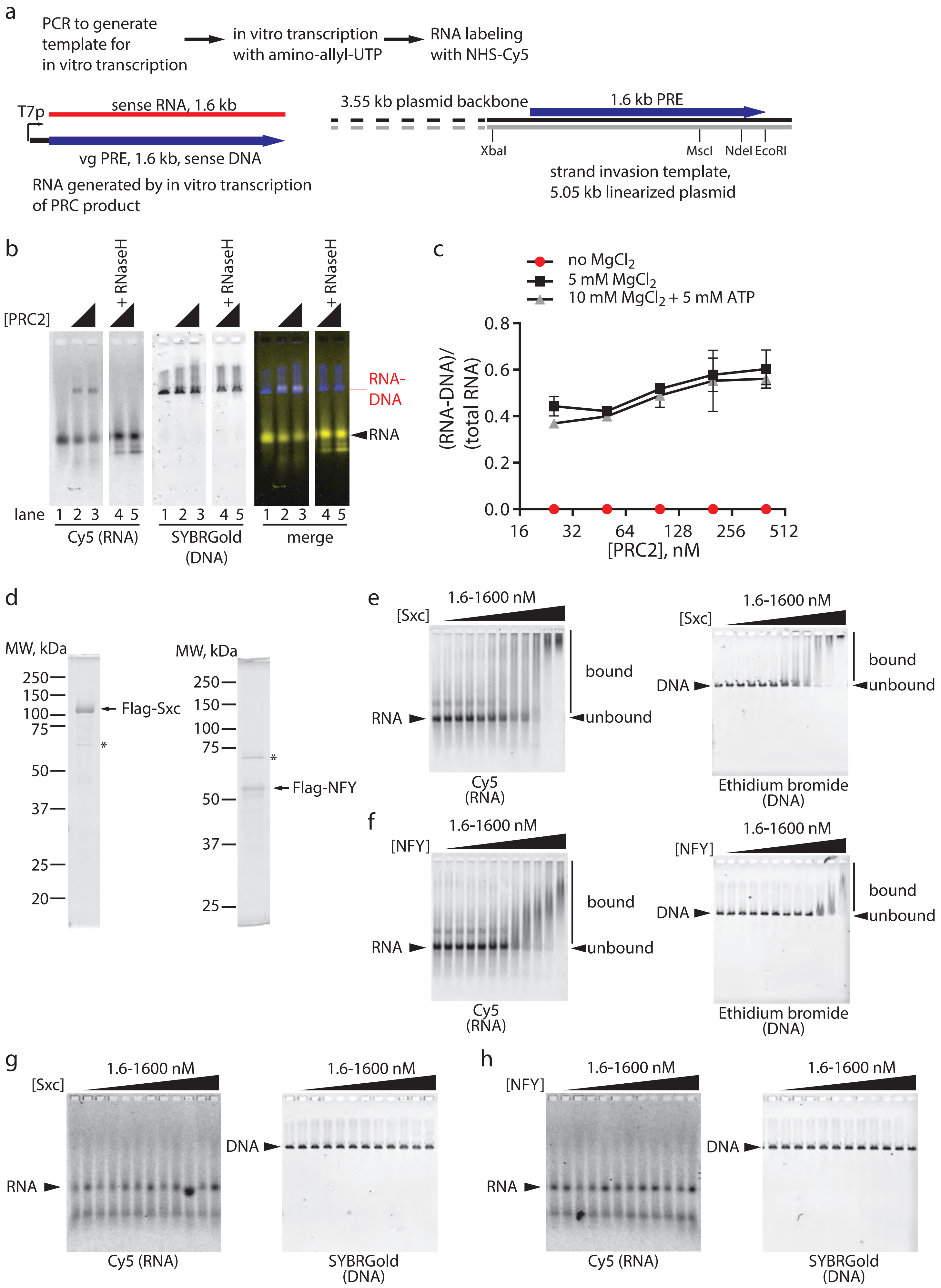


Alecki et al., Extended Data Fig. 6

**Extended Data Fig. 7 Co-fractionation of strand invasion activity with PRC2 through size exclusion chromatography.** a. 45 µg of PRC2-Flag-Su(Z)12) were fractionated on a Superdex 200 PC3.2/300 column. b. SDS-PAGE (10%) of fractions from size exclusion column. 7 µl of each fraction were loaded on the gel, which was stained with SYPRO Ruby. c. Strand invasion assay of the size column fractions. 4 µl of each fraction was used in this assay with 0.3 nM DNA and 0.18 nM RNA. Similar co-fractionation of activity with PRC2 was observed with two additional preparations of PRC2 using glycerol gradient sedimentation, and a second PRC2 preparation using size exclusion chromatography.

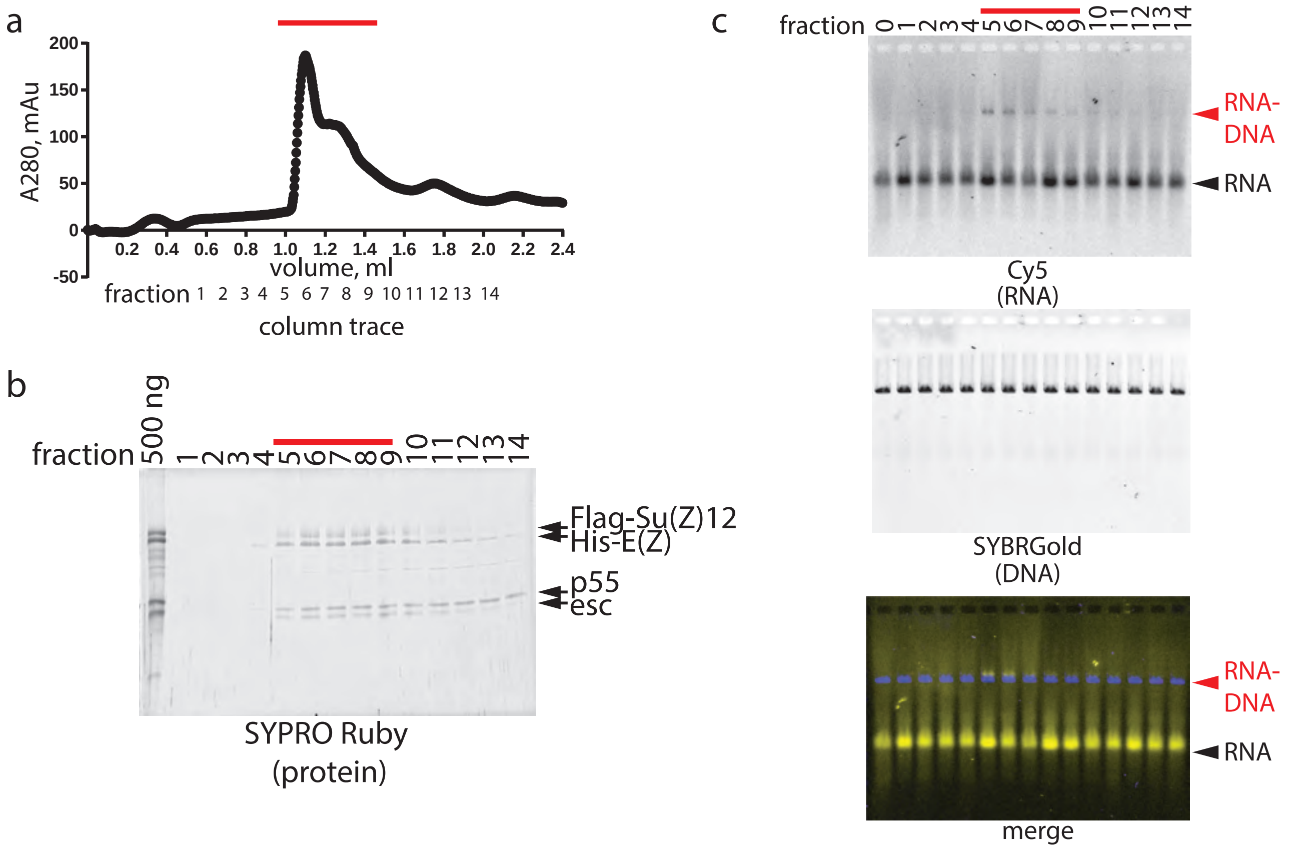


Alecki et al., Extended Data Fig. 7

**Extended Data Fig. 8 Full gels of assays shown in Fig. 4 and confirmation of RNA-DNA hybrids induced by hPRC2.** a. Gels from Fig. 4a (time course). b. Gels from Fig. 4c (order of addition of DNA and RNA). d. Gels from Fig. 4e (RNA titration). d. Gels from Fig. 4g (DNA titration). e. Strand invasion products are sensitive to RNaseH (lanes 5, 6), but resistant to RNaseA (50 pg) (lanes 8, 9). For this analysis, strand invasion products were purified by phenol-chloroform extraction followed by ethanol precipitation. Under these conditions, faint bands corresponding to strand invasion products are observed for hPRC2-EZH2, which were not observed in our standard reactions (Fig. 4i).

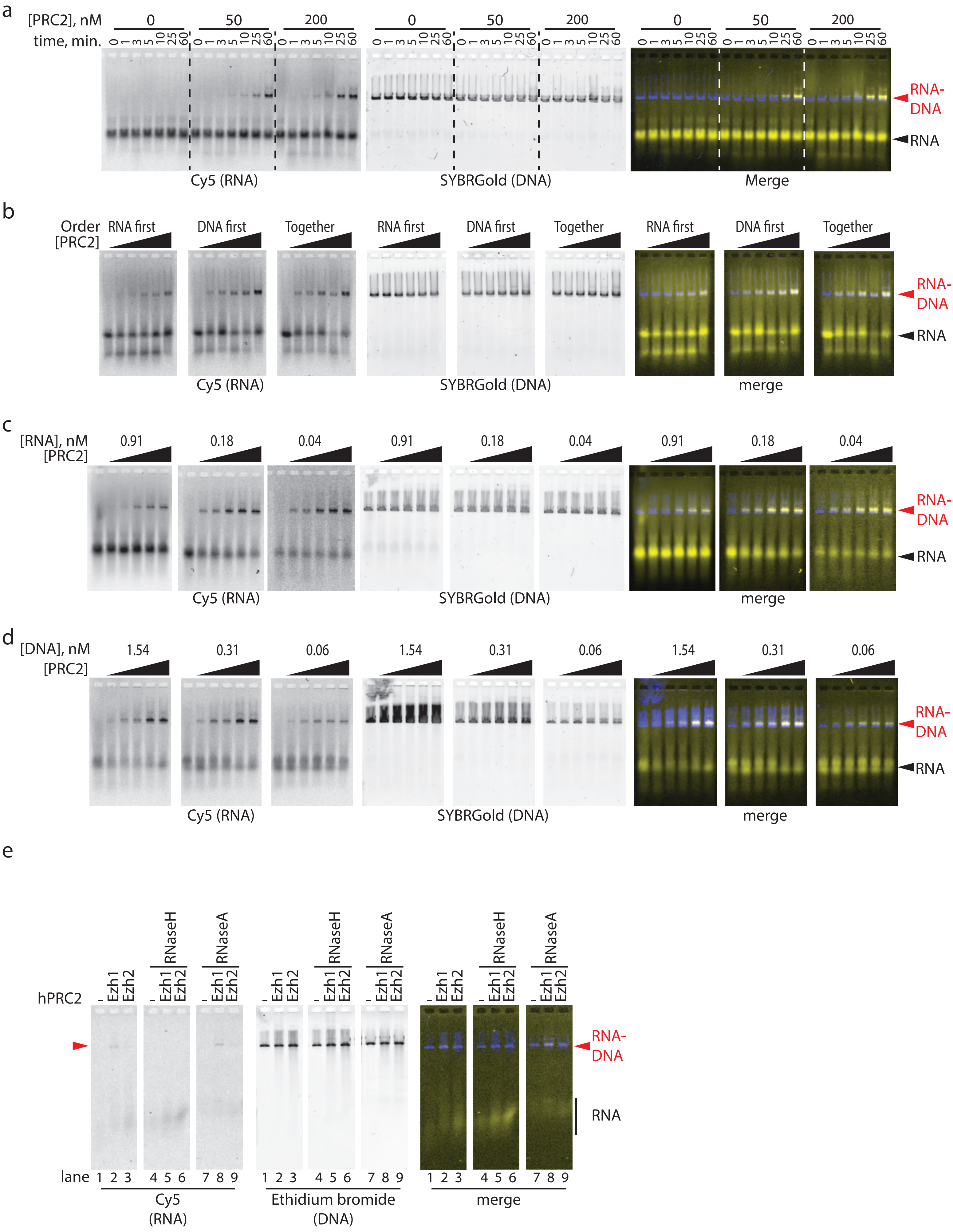


Alecki et al., Extended Data Fig. 8

**Extended Data Fig. 9 Models for strand invasion mediated by PRC2** Three hypothetical schemes for the strand invasion reaction. a. step 1: PRC2 binds DNA; step 2: formation of "open" DNA complex; step 3: formation of ternary complex DNA-PRC2-RNA; step 4: strand invasion. As indicated free RNA is expected to compete for PRC2-DNA interactions. For step 3, two possible scenarios are depicted. In the top scheme, RNA enters the complex with a second PRC2, consistent with PRC2 dimerization (note the dimer could form during the reaction, as shown here, or a preformed dimer bound to DNA could recruit the RNA).In the bottom scheme, RNA enters the complex via a second binding site on PRC2. We do not favour this possibility because biochemical evidence argues against two nucleic acid binding sites in the complex [22]. In the 4$^{th}$ step, strand invasion occurs, and PRC2 is released. b. Steps 1 and 2 are as in A; step 3: RNA pairs with the open DNA leading to strand invasion and possibly displacing PRC2 (step 4). c. Classical "inverse strand invasion" mechanism in which a PRC2-DNA filament is formed which allows RNA strand invasion. We do not know if PRC2 forms such filaments, but the stoichiometry of PRC2:DNA used in our reactions (1:62bp-1:4bp) is consistent with this possibility [34]. We note that these models are very general and other schemes are possible.

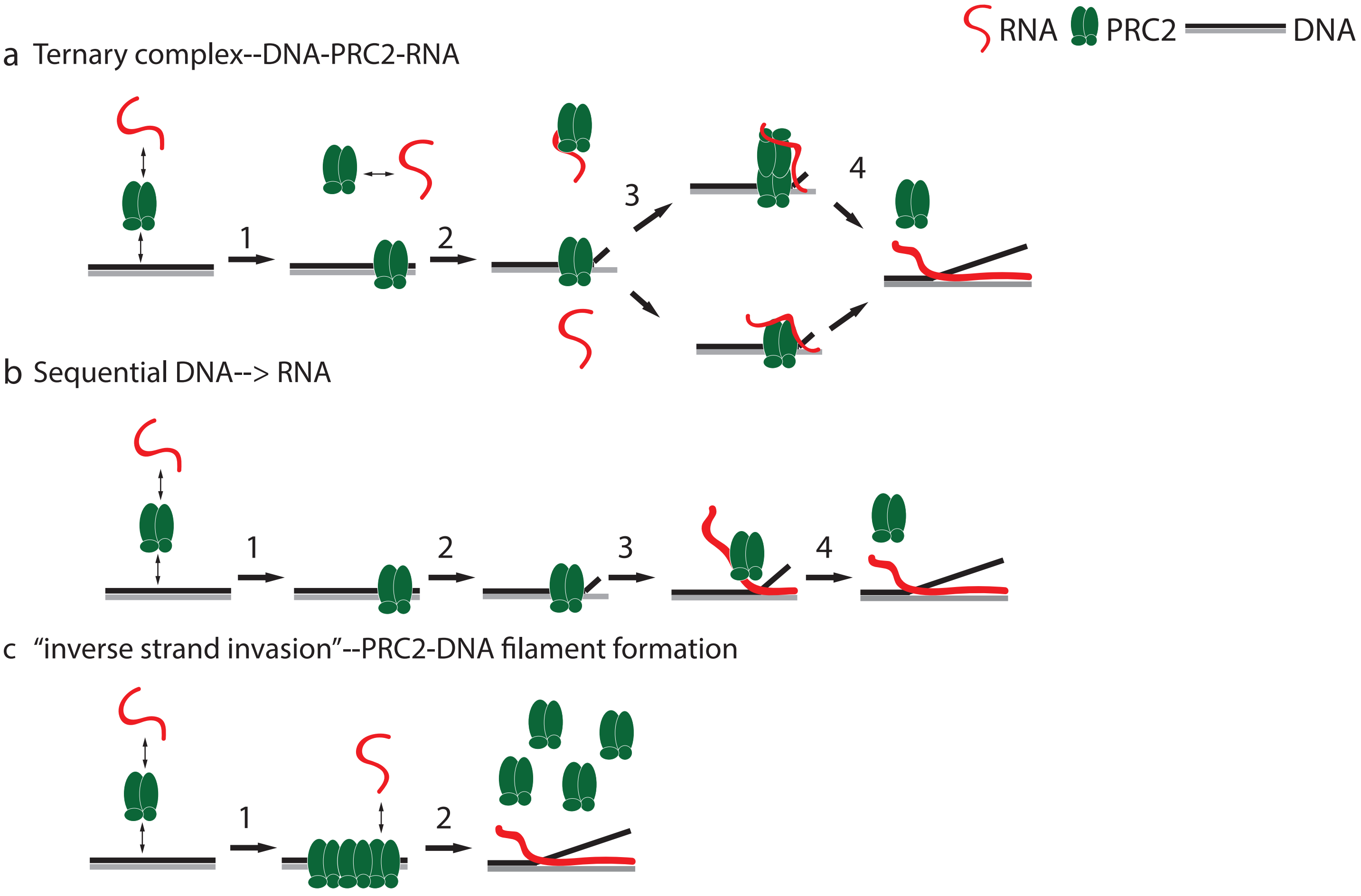
RNA
PRC2
DNA
a Ternary complex--DNA-PRC2-RNA
1
2
3
4
b Sequential DNA--> RNA
1
2
3
4
c "inverse strand invasion"--PRC2-DNA filament formation
1
2

**Extended Data Fig. 10 Model for the role of R-loop formation driven by PRC2 in PcG gene silencing.** 1) Widely expressed PRE binding transcription factors (TFs) target PRC1 and PRC2 PREs. 2) If RNA is produced at the PRE, it may bind PRC2. 3) PRC2 mediates strand invasion, forming an R-loop with part or all of the RNA. Although our strand invasion assay uses linear DNA templates, we anticipate that opening of the DNA, but not a DNA end, would be required at PREs. DNA opening could occur through transcription, or involve topoisomerases. Topo II interacts genetically with PcG genes and co-localizes to PRES. 4) R-loop formation may release PRC2 from the RNA, allowing it to bind to chromatin. The R-loop will also synergize with TFs that recruit PRC1 to increase its binding to the PRE. 5) PRC1 and PRC2 modify surrounding chromatin to repress transcription. It is also possible (although not shown), that R-loops interfere with transcription directly, or prevent binding of transcription activators. We have not drawn any TFs binding to R-loops, although some DNA binding proteins may bind RNA-DNA hybrids [37] or the displaced single-stranded DNA [38]. For simplicity, we have drawn persistent R-loops, but it is likely that R-loops are dynamic through the cell cycle. R-loops may also not be persistently required. For instance, we do not know if the final repressed chromatin structure includes an R-loop or if R-loop formation and chromatin structure changes are sequential.

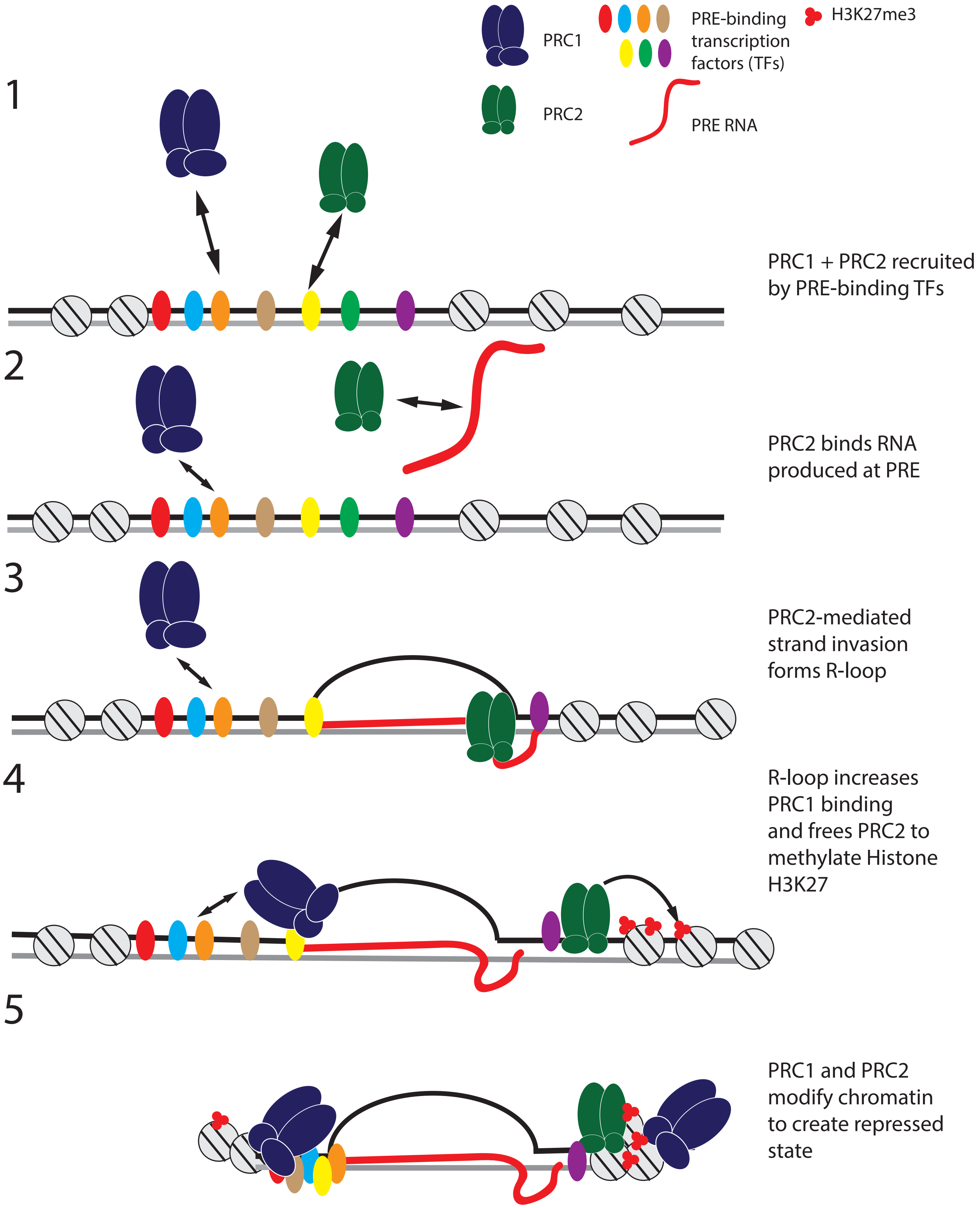


Alecki et al., Extended Data Fig. 10
